# The urge to blink in Tourette syndrome

**DOI:** 10.1101/477372

**Authors:** Haley E. Botteron, Cheryl A. Richards, Tomoyuki Nishino, Keisuke Ueda, Haley K. Acevedo, Jonathan M. Koller, Kevin J. Black

## Abstract

Functional neuroimaging studies have attempted to explore brain activity that occurs with tic occurrence in subjects with Tourette syndrome (TS). However, they are limited by the difficulty of disambiguating brain activity required to perform a tic, or activity caused by the tic, from brain activity that generates a tic. Inhibiting the urge to tic is important to patients’ experience of tics and we hypothesize that inhibition of a compelling motor response to a natural urge will differ in TS subjects compared to controls. This study examines the urge to blink, which shares many similarities to premonitory urges to tic. Previous neuroimaging studies with the same hypothesis have used a one-size-fits-all approach to extract brain signal putatively linked to the urge to blink. We aimed to create a subject-specific and blink-timing-specific pathophysiological model, derived from out-of-scanner blink suppression trials, to eventually better interpret blink suppression fMRI data. Eye closure and continuously self-reported discomfort were reported during five blink suppression trials in 30 adult volunteers, 15 with a chronic tic disorder. For each subject, data from four of the trials were used with an empirical mathematical model to predict discomfort from eye closure observed during the remaining trial. The blink timing model of discomfort during blink suppression predicted observed discomfort much better than previously applied models. However, so did a model that simply reflected the mean time-discomfort curves from each subject’s other trials. The TS group blinked more than twice as often during the blink suppression block, and reported higher baseline discomfort, smaller excursion from baseline to peak discomfort during the blink suppression block, and slower return of discomfort to baseline during the recovery block. Combining this approach with observed eye closure during fMRI blink suppression trials should therefore extract brain signal more tightly linked to the urge to blink.

## 1. Introduction

> *Often the effort to control these wild sensations seems to be more than the human spirit can bear. … At the onset of an impulse … it does not seem possible for the state to be relieved by anything other than the action in progress.*
>
> — — (Bliss, Cohen, & Freedman, 1980), discussing premonitory urges to tic

Tourette syndrome (TS) is a chronic neuropsychiatric developmental disorder characterized by motor and vocal tics beginning in childhood (American Psychiatric Association, 2013). Tics differ from other abnormal movements because they can be suppressed for some period of time, although suppression is associated with increasing discomfort (Jankovic, 1997). Behavior therapies are first-line treatments for TS and focus on the link between tics and their preceding urges. Inhibition of tics is part of daily life for most people with TS and it is also an important part of the theory and technique of exposure and response prevention (Verdellen, Keijsers, Cath, & Hoogduin, 2004; Woods et al., 2008). Previous functional imaging studies attempting to explore brain activity involved with tic suppression have been complicated by the fact that successful suppression is unavoidably accompanied by a decrease in movement, which reflects inverse changes in brain activity related to tics during suppression.

Over 90% of people with tics report premonitory urges, meaning an uncomfortable sensation or the perceived need to perform the tic (Kwak, Dat, & Jankovic, 2003; Reese et al., 2014; Crossley & Cavanna, 2013; Miguel et al., 2000; Cavanna, Black, Hallett, & Voon, 2017). Such urges build up prior to a tic and then are transiently relieved by the tic before building up again (Leckman, Walker, & Cohen, 1993; Reese et al., 2014). In fact, many patients with TS feel that the premonitory phenomena are primary, rather than the tic *per se* (Bliss, Cohen, & Freedman, 1980; Bullen & Hemsley, 1983). This view contributes to the interpretation that tics may be maintained and reinforced by the consequent reduction in discomfort (premonitory urge); premonitory urges thus “may represent an important enduring etiological consideration in the development and maintenance of tic disorders” (Specht et al., 2013). Supporting evidence for this view includes the observation that patients who believe tics are unavoidable or barely suppressible once they experience a premonitory urge have higher impairment ratings on the YGTSS and higher premonitory urge severity (Steinberg et al., 2013). A tic—especially one associated with a premonitory urge—may therefore be viewed as a transient failure of motor inhibition.

The urge to tic shares many characteristics with natural urges, such as the urge to blink after keeping one’s eyes open for an extended period of time (Jackson, Parkinson, Kim, Schüermann, & Eickhoff, 2011; Belluscio, Tinaz, & Hallett, 2011). The present report was motivated by a study intended to examine the brain activity correlated with the urge to blink, or with imminent failure of blink inhibition, in TS and control subjects. Importantly, the urge to blink can be examined in subjects with or without tics. A necessary early step in such an effort is to identify moments during which the urge to blink fluctuates. Previous functional MRI (fMRI) studies examining the urge to blink associated with voluntary eye blink suppression used two approaches to do so. The simplest approach compared mean BOLD signal during blink suppression blocks to mean signal during blocks in which subjects blinked normally, represented by the square-shaped (red) function in Figure 1A (Lerner et al., 2009; Mazzone et al., 2010). Berman et al. hypothesized that relevant brain activity might be identified better by a linear model of urge build-up during blink suppression, as in Figure 1B (Berman, Horovitz, Morel, & Hallett, 2012). In fact, their study reported urge severity on a group level that somewhat resembled this sawtooth model.

**Figure 1.**
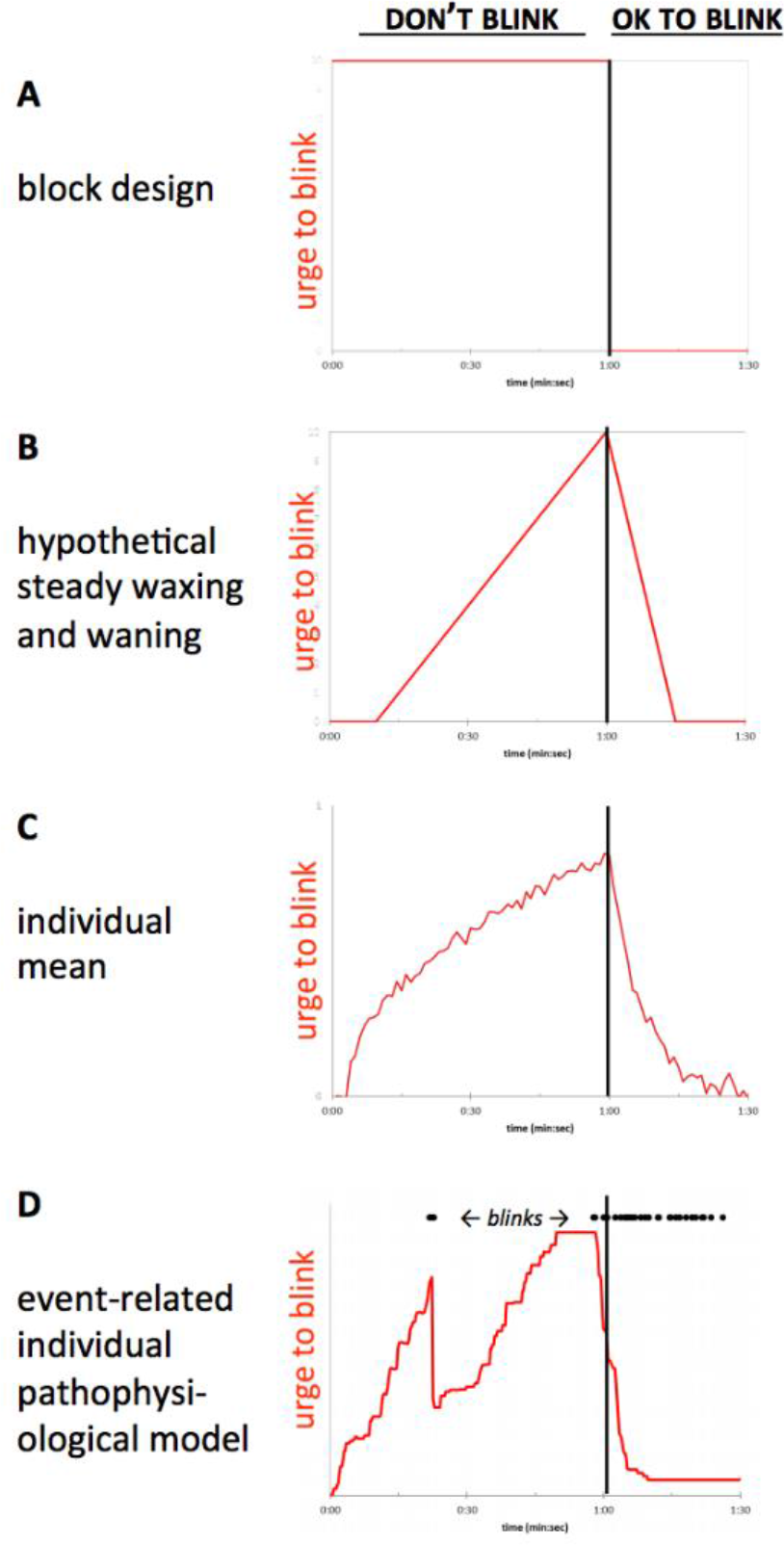
(A) Block design comparing “OK to blink” and “don’t blink” blocks (Mazzone et al., 2010). (B) The sawtooth approximation of urge to blink used by Berman and colleagues (Berman, Horovitz, Morel, & Hallett, 2012). (C) An example average of self-rated time–urge curves from all of a given subject’s trials. (D) An event-related, individualized model proposed for this study, in which the time–urge curve depends on the specific timing of the blinks observed in a single trial, with urge to blink decreasing after each eye closure. In this model, a given time–urge curve would thus apply only to a single experimental block in a single subject. The curves in (C) and (D) were drawn for illustration, without reference to data.

These models enabled analysis of fMRI blink suppression data, but we saw potential for improvement in trying to achieve the same goal. First, these block and sawtooth patterns of discomfort or urge to blink during blink suppression were hypothesized rather than derived from subject-reported data. We found early on that self-reported discomfort during blink suppression followed neither the box nor the sawtooth model (Claudio Torres, Black, Richards, & Black, 2014). Second, urge ratings over time may vary by subject. Figure 1C represents a simple average of several self-report trials from one subject, an approach that would address these two concerns. Third, neither model takes into account the likely effects on urge from any eye closures that occur during a blink suppression block. Such effects would depend on the timing of blinks observed in a specific block; at the extreme, if a subject continued blinking at his/her baseline rate when instructed not to blink, one would expect no buildup in that subject’s urge to blink. Thus we developed a model to predict urge to blink based on observed blink timing, expecting that the urge to blink would decrease somewhat after each period of eye closure, as diagrammed in Fig. 1D. We hypothesized that such a model, taking into account observed eye closures, would better fit subjects’ reported discomfort when instructed not to blink.

Our objectives with this study were:

1. To create a subject-specific and blink-timing-specific model from out-of-scanner blink suppression trials in the same subject (as in Fig. 1D), to facilitate eventual application to identify brain regions with discomfort-related activity during a “don’t blink” fMRI session.
2. To test whether the new model more accurately predicted discomfort based on blink timing than did the square or sawtooth models shown in Fig. 1 A,B, or the individual mean model in Fig. 1C.
3. To estimate the accuracy of the new model compared to direct subject report, using a leave-one-out cross-validation (LOOCV) approach.

## 2. Materials & Methods

### 2.1 Ethics approvals

This research was approved in advance by the Washington University Human Research Protection Office (protocols #201303016 and #201506018). All subjects participated only after written informed consent.

### 2.2 Subjects

We examined data from two studies.

**Study 1.** This study included 18 adult volunteers who continuously reported urge to blink during blink suppression; these data were examined to improve our methods for measuring urge to blink, and to select models to analyze future data. We had hypothesized that effort or ability to suppress blinks might vary independently of discomfort: One person might have relatively high discomfort while holding the eyes open, but due to greater effort or greater inhibitory ability might blink no more than another person who experienced relatively little discomfort. Thus we chose to separate urge to blink into two hypothesized components, discomfort and effort, and track each separately. However, in our initial pilot data, mean self-rated discomfort and effort curves during the blink suppression phase were fairly similar (Claudio Torres, Black, Richards, & Black, 2014), so the remainder of this report focuses on discomfort.

**Study 2.** This study provided the data reported below. It included 15 subjects with TS (age 28 ± 4.2 years, range 18-30; 6 Male) and 15 control subjects (age 26 ± 2.2 SD; 6 Male). Eligible participants for the TS group included those who met the criteria for DSM‑5 Tourette’s Disorder or Persistent Motor or Vocal Tic Disorder (American Psychiatric Association 2013) with tics in the past week; they also must have had a tic involving the right eyebrow, eyelid or cheek at some point. Nine subjects had OCD and four had ADHD. Exclusion criteria for all subjects included contraindication to MRI, intellectual disability, substantial autistic traits (SRS T score ≥ 75; (Constantino, 2013)), dementia, currently active major depression or substance abuse, a neurological or general medical condition that would interfere with study participation, or lifetime history of psychosis, mania or somatization disorder. Potential controls were also excluded for a past or present tic disorder in the subject or a first-degree relative. TS symptom severity was assessed using the Yale Global Tic Severity Scale (YGTSS; (Leckman et al., 1989)) and TS symptom history using the Diagnostic Confidence Index (DCI; (Robertson et al., 1999)). The overall severity of premonitory urges was assessed using the Premonitory Urge for Tics Scale (PUTS; (Woods, Piacentini, Himle, & Chang, 2005)), and each subject also completed the Beliefs About Tics Scale (BATS; (Steinberg et al., 2013)). Symptoms of OCD and ADHD were rated using the Yale-Brown Obsessive Compulsive Disorder Scale (Y-BOCS; (Goodman, 1989)) and ADHD Rating Scale (Barkley, 1990) respectively.

### 2.3 Data collection

Demographic data were collected and managed using REDCap electronic data capture tools hosted at Washington University in St. Louis (Harris et al., 2009). Each subject was observed for a two-minute baseline period to measure the baseline blink rate, and then performed two blink suppression tasks. In each of these tasks, the subject performed 5 trials seated in front of a computer monitor, each characterized by 60 seconds of blink suppression (the monitor read “DON’T BLINK”), followed by 30 seconds during which they were allowed to blink freely (“OK TO BLINK”). During one task (“discomfort trials”), each subject continuously rated discomfort using the vertical position of a mouse-controlled pointer, with continuous feedback that interpreted the position on a 0-9 scale (Subjective Units of Distress) (Benjamin et al., 2010; Specht et al., 2013). The other task (“effort trials”) was identical except that, rather than rating discomfort, subjects were instructed to rate the effort being used to keep the eyes open. Order of tasks (discomfort, effort) was balanced across subjects. The baseline blinking condition was included to ensure that subjects exhibited relative suppression during the “don’t blink” task. Eye closures were recorded using a video camera. Tic subjects were also observed by video recording on the screening day for 5 minutes of baseline (“OK to tic”) and 5 minutes of tic suppression.

#### 2.3.1 Blink detection

Blinks from the first two seconds of each trial were ignored to allow for the participant to adjust to the start of each condition. Because many subjects used only a subset of the full range from 0-9, the discomfort scores were *z*-standardized for each subject to reduce variance across subjects, as suggested by (Brandt et al., 2016): *z* = (*x*_*i*_ − *x̅*)/*SD*. The SD was computed over all ratings recorded by the subject in all blocks. A custom program written in C++ was modified from code kindly provided by Dr. Aleksandra Królak to identify whether on each frame of the video the subject’s eyes were closed. The new program compares the intensity histogram of a region-of-interest (ROI) centered on each iris in each video frame to a baseline frame in which the eyes were open. The ROIs had been marked manually on the baseline frame. The ROIs of the subsequent frames were centered automatically using a motion detection algorithm to follow head movement. The root mean squared error (RMSE) of the ROI intensity histogram comparison was computed for each frame, and frames in which the RMSE value exceeded a specified threshold were interpreted as eye closures. The results were then visually checked to see if the program happened to miss a blink due to head movement and timing was adjusted as needed. A Python script was used to eliminate suprathreshold deviations lasting less than 100 ms and to create a binary time series indicating eye closure at each video frame. The eye closure time series and discomfort ratings were synchronized and resampled to identical .25 s blocks. For each quarter-second time bin, the eye closure fraction *f*_*i*_ was defined as the number of eyes-closed frames within a given bin, divided by the total number of frames in that bin. Blink rate appeared similar over repeated trials, so means across trials were compared. Several subjects were outliers, so blink rates were compared between groups using the nonparametric Mann-Whitney test.

#### 2.3.2 Tic detection

To analyze tic suppression, the videos were clipped and renamed so that the rater was blind to the condition. Author KU reviewed these recordings and indicated the presence of a tic with a button press using a slightly modified version of our TicTimer software (Black, Koller, & Black, 2017). Tic frequency and the number of 10-second tic free intervals per minute were calculated from the TicTimer output and tic suppression was calculated using the difference between the two conditions as a fraction of the baseline.

### 2.4 Model development

We examined data from Study 1 and Study 2 to choose the form of an empiric mathematical model of self-reported discomfort. In order to build a subject-specific and blink-timing-specific model we first looked at subjects’ raw discomfort scores and settled on a model in which discomfort during the “don’t blink” block increases as the square root of time, with brief dips in discomfort after eye closure, and discomfort during the “OK to blink” block decreases exponentially back to its baseline level when the eyes are closed. Details on how the model was created are provided in the Appendix. We also compared these results to a simpler “individual mean model” comprising the average across trials of a subject’s z-scored discomfort ratings.

#### 2.4.1 Bayesian parameter estimation

The parameters *a*, *b*, *c*, *h*, *k* and *r* for the blink timing model were fit to self-reported discomfort over time using observed blink timing and the Bayesian Data-Analysis Toolbox, which implements a Markov chain Monte Carlo method (Bretthorst & Marutyan, 2016; Bretthorst, 1988). Prior probabilities for each parameter were chosen as follows. To minimize the risk that the priors would constrain the estimation, we assumed uniform priors over [0, 10^4^] for *a*, [−10, 10] (standard deviation units) for *b*, [0,30] seconds for c, and [0,10] s^−1^ for r. The data presented in the appendix suggested the following priors for the coordinates (*h*,*k*) of the vertex of the parabola representing a transient decrease in discomfort after eye closure: for *h*, a uniform prior from 0 to 10s, and for *k*, a Gaussian prior with mean −0.4, S.D. 1.0, range [−1,0].

#### 2.4.2 Accuracy

To evaluate the accuracy of a model’s prediction of discomfort, we used data from 4 trials to predict discomfort of the 5th trial, in a leave-one-out cross-validation (LOOCV) approach. Specifically, we excluded one of the 5 suppression trials and estimated model parameters from the data from the remaining 4 trials, using the “Enter ASCII Model” package from the Bayesian Data-Analysis Toolbox. If the model served as a good predictor, then the discomfort predicted by that model with parameters based on the remaining trials should closely match the actual discomfort reported during the excluded trial. For the blink timing model, the parameters based on the remaining trials were applied to the observed blink timing of the excluded trial to estimate discomfort for the excluded trial. Each subject had 5 complete suppression trials, and therefore 5 sets of estimated parameters were computed.

### 2.5 Comparison to other models

To compare the 4 models, we calculated the correlation values for each LOOCV step, *i.e.* the Pearson *r* value comparing the model fit to the remaining 4 trials *vs.* the data from the trial left out. Correlation was used to measure model accuracy because the individual subject time–urge signal, predicted from blink timing in the fMRI session and the parameters identified from the out-of-scanner blink suppression trials, will be used as a contrast in a general linear model to identify regions in each subject that correspond to subject discomfort during periods of effective blink suppression in the scanner. Pearson *r* values were compared between models using the t-test after Fisher’s *z* transform. The models were also compared using the binomial test in terms of how often one model outperformed another, with the null hypothesis that either model is equally likely to be superior on each run or in each subject.

## 3. Results

A total of 28 participants completed the blink suppression trials to be analyzed in the results. Data were excluded from one control subject who appeared not to understand the task and did not suppress blinking in the blink suppression condition as compared to the baseline condition. Another participant was excluded from the model comparison analysis because the first 30 seconds of the first task trial was not recorded on video. All participants rated discomfort and effort on separate trials.

### 3.1 Blink rate

The median number of blinks during a 60-s suppression trial was 2.0 for controls (interquartile range 0.8, 3.0) and 4.6 (3.1, 9.1) for TS subjects. During the 30 seconds in which they were told it was OK to blink, control subjects showed a median of 47.2 blinks per minute (40.0, 72.4) and TS subjects 46.4 (41.6, 54.4) blinks per minute. The groups differed significantly on blinking during the suppression condition (W = 42, p = 0.012) but not in the OK-to-blink period (W = 106, p = 0.712). The groups did not differ significantly on blink rates during the initial 2-minute baseline period (W = 77, p = 0.357), with baseline rates of 17.5 (11.5, 21.5) blinks per minute for controls and 25.5 (15.8, 31.5) for the TS group. We also computed a secondary measure, blink suppression efficacy, as a fraction of the baseline rate. Blink suppression efficacy was 91.6% (85.1%, 95.6%) for the control group and 79.2% (71.4%, 86.4%) for the TS group (W = 52, p = 0.038). In the TS group, the minimum blink suppression efficacy was 52%.

### 3.2 Tic suppression

By contrast, tic suppression efficacy in the TS group was less complete and more variable; median tic suppression was only 56% (range −29% to 100%). Blink suppression and tic suppression within TS subjects were not correlated (r = −0.13, p = 0.68).

### 3.3 Discomfort ratings

The raw discomfort ratings averaged across all trials for each subject did not differ significantly between groups (TS 3.2 ± 1.7, control 3.6 ± 1.0, p = 0.50). However, scaled discomfort ratings and blink timing varied substantially among subjects, supporting the individual-specific pathophysiological modeling approach. The model represented by Equations 3 and 5 of the appendix was applied to each subject and the average parameter estimates are summarized in Table 1.

**Table 1.** Comparison of average parameter values for subjects with TS and control participants. *: values are mean ± SD, except for model fit, for which the t test was applied to Fisher’s z(r) scores; the mean ± SD for each group were converted back to r values and reported here as r(mean) (r(mean−SD)–r(mean+SD)). † SUDS = Z scores for subjective units of distress.

| Block | Model parameter | Units | TS, N=14 * | Ctrl, N=14 * | p (2-tailed t test) |
| --- | --- | --- | --- | --- | --- |
| both blocks | model fit (Pearson's $r$) | — | 0.82 (0.59–0.93) | 0.9 (0.79–0.95) | $3.36 \times 10^{-5}$ |
| don't blink | a (change with time in discomfort) | SUDS $\cdot s^{-1/2}$<br>† | $0.27 \pm 0.1$ | $0.33 \pm 0.05$ | $3.49 \times 10^{-4}$ |
| don't blink | b (baseline discomfort) | SUDS | $-0.879 \pm 0.29$ | $-1.06 \pm 0.23$ | $2.13 \times 10^{-4}$ |
| don't blink | c (time offset of change in discomfort) | s | $6.53 \pm 9.58$ | $6.41 \pm 5.93$ | 0.93 |
| don't blink | h (half duration of reduction with one blink) | s | $4.7 \pm 3.22$ | $4.89 \pm 4.09$ | 0.76 |
| don't blink | k (amplitude of reduction with one blink) | SUDS | $-0.31 \pm 0.18$ | $-0.34 \pm 0.28$ | 0.41 |
| OK to blink | r (recovery rate) | $s^{-1}$ | $1.72 \pm 1.4$ | $2.49 \pm 2.57$ | 0.03 |

The differences in the mean parameter values for *a*, *b* and *r* can be understood by their effect on the time–discomfort curve estimated for blink timing from one trial, as shown in Fig. 2.

**Figure 2.**
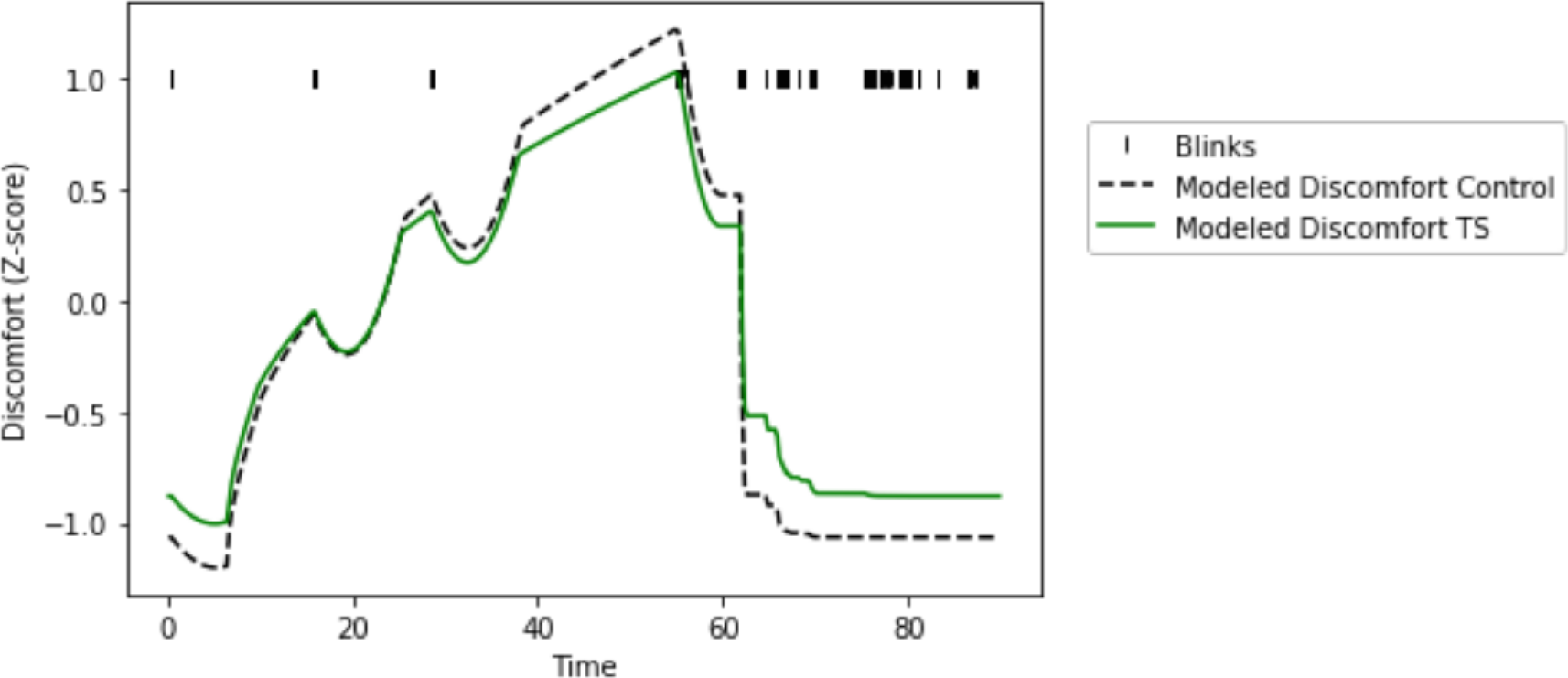
Interpreting the different parameter values between TS and control subjects. Discomfort during a suppression trial modeled using the average parameter values from Table 1 for both TS and control subjects, applied to a single set of observed blinks. The timing of blinks is shown by vertical black marks near the top of the graph, to help visualize the change in discomfort after a blink. Longer eye closures appear as thicker black marks.

### 3.4 Model comparison

In the LOOCV testing, the model that returned the highest correlation *r* with the reported discomfort was the individual mean model for 13 subjects, the blink timing model for 13 subjects, the sawtooth model for 2 subjects, and the box model for none. If all models were equally likely, the chance that any model would “win” in 13 or more subjects is .011 (binomial test). The correlations differed significantly (ANCOVA on mean *z*(*r*) values for each subject, with factors for model, diagnosis and their interaction; model significant at p<.001, diagnosis not significant, interaction p=0.08). In post hoc tests (Tukey HSD), the box model performed worse than each of the others (p<.001 for each), while the individual mean (p=.029) and blink timing models (p=.005) were each superior to the sawtooth model but did not differ significantly from each other (p=.929).

## 4. Discussion

This report describes novel methods to impute momentary discomfort associated with the urge to blink during blink suppression, based on the actual observed timing of blinks and the individual subject’s rating of discomfort. We show that both new methods more accurately predict discomfort than simpler models that do not account for individual subject sensitivity nor the timing of blinks. This result raises our expectation of finding regions with a pertinent BOLD signal when individual response characteristics are taken into account, but of course, that expectation remains to be tested.

Our data provide some interesting observations beyond these methodological results. The 15-fold ratio of the mean blink rates during the blink suppression and recovery phases indicates that overall, subjects successfully suppressed the urge to blink. However, motor response differed between groups. Subjects with TS averaged an additional 2.6 blinks during each suppression task than did controls, who often went the whole 60 seconds without blinking. By contrast, the number of blinks between the two groups during the rest period did not differ significantly. The group difference in blink suppression is consistent with the hypothesis that people with TS may exhibit a motor inhibition deficit compared to controls when engaged in real-world suppression of an uncomfortable urge to move. This finding is interesting given that deficits in inhibition have long been hypothesized in TS but have not been reliably identified in most standard psychological tests (Morand-Beaulieu et al., 2017). Conceivably, inhibiting longstanding (overlearned) motor responses to sensory discomfort differs in important ways from inhibiting button presses as directed by an experimenter (Jackson, Parkinson, Kim, Schüermann, & Eickhoff, 2011). While tic subjects were less effective at suppressing blinks than controls, the correlation between blink suppression and tic suppression within TS subjects was not significant. This result may suggest that although blink suppression and tic suppression both reflect somatomotor inhibition, the two may not overlap completely.

Additionally, subjects in the TS group experienced discomfort from blink suppression in a quantitatively different manner than did controls. Specifically, in the blink timing model, the TS group had higher baseline discomfort, smaller excursion from baseline to peak during the blink suppression block, and slower return to baseline during the recovery block (see Table 1 and Fig. 2). However, these differences come from the scaled data, and raw baseline discomfort scores were not higher in the TS group. This may reflect the fact that reported discomfort (both within and across subjects) was also more variable in the TS group, as reflected by the quantitative model fit to the data. Thus these group differences in reported discomfort should be interpreted with caution.

### 4.1 Comparisons

While a previous study by (Berman, Horovitz, Morel, & Hallett, 2012) examined the fMRI data for a signal that matched a hypothesized urge model or mean urge across subjects, our analysis can account for responses to the timing of each subject’s blinks in the scanner and his or her individual response characteristics. Berman and colleagues did perform an analysis of their BOLD data on an individual subject basis, but their approach identified signal temporally coinciding with blinks, not urge ((Berman, Horovitz, Morel, & Hallett, 2012); Berman personal correspondence to KJB, 2/21/2018). Our subject-specific and blink-dependent models had a significantly better correlation to urge than did the two simpler models proposed by Berman, Mazzone, and colleagues.

Brandt and colleagues (2016) also examined the timing of urge and related movement to tics in subjects with TS. However, they only used blink timing and suppression with controls, and tic timing and suppression with TS subjects, while we aimed to compare the same task (blink suppression) across groups. Our results supported their finding that an increase in urge intensity is temporally related to blinks, increasing until a blink is executed and immediately decreasing afterward. Another interesting result from their study was that all their healthy control subjects, but only two with TS, reported an increase in their self-reported experience of urge intensity when they were required to monitor and rate their current urge. We asked subjects to rate urge intensity continuously, so we cannot verify that finding, but in future studies it may be interesting to investigate the subject’s attention to urge, as one interpretation may be that patients with TS pay more attention to their urges in daily life. However, tic frequency in TS decreases with self-monitoring of tic and urge occurrence (King, Scahill, Findley, & Cohen, 1999; Misirlisoy et al., 2015), though by contrast, watching oneself tic increases tic frequency (Brandt, Lynn, Obst, Brass, & Münchau, 2015). It is possible that our subjects may have heightened discomfort ratings because of increased attention to their discomfort, but this result would be carried through all trials and thus should not confound the comparisons reported above.

In perhaps the first attempt to image neural correlates of the urge to blink, Sandor and colleagues reported pilot data on two other approaches to elicit urge to blink in the scanner. One focused on the few seconds just before the first blink versus the few seconds after a 5-s eyes closed rest condition, and the other compared cued blinking at twice (control) or half (urge) the subject’s observed baseline blink rate (Sandor, Silveira, Mikulis, & Crawley, 1998). A block design was used to analyze each comparison. More recently, Abi-Jaoude *et al.* used fMRI to study increased failure of blink suppression in healthy subjects over repeated trials (Abi-Jaoude, Segura, Cho, Crawley, & Sandor, 2018). Although their study design differed somewhat from ours, we did not see in either the TS or control group the increase in escape blink frequency over repeated trials that formed the basis of their analyses.

### 4.2 Limitations

This study relies on self-report for discomfort, a subjective experience that people may report differently. However, we believe this is a reasonable proxy and will apply it to BOLD signal from the fMRI data described above with the goal of identifying a rater-independent, objective, neuroimaging measure of discomfort. This study did not exclude potentially confounding factors such as OCD and ADHD. Their prevalence in TS makes exclusion difficult in a study of this size. Another potential limitation is the use of discomfort as a proxy for urge. We used discomfort because it seemed an easier term to understand and rate than “urge,” and the urge to blink, one would think, would correspond very closely to discomfort. We had wanted to separate urge into two dimensions, discomfort and effort; however, these did not differ significantly in our initial pilot data (Claudio Torres, Black, Richards, & Black, 2014). As noted above, group differences on the scaled discomfort ratings should be interpreted with caution. Finally, any study of blinking in tic patients raises the question of whether blinking tics affect the analysis. However, in this sample only three of the subjects with TS reported blinking tics in the past week or had a blinking tic observed on exam. Furthermore, the two groups did not differ in terms of blink rate during the recovery period. If blinking tics were prominent enough to overcome blink suppression, one would expect that they would also increase the number of blinks when the subjects were free to blink during the ‘OK to blink’ period. The most likely explanation for these observations is that blinking tics did not substantially affect our results.

### 4.3 Future directions

The next step will be to apply this model to fMRI data from the same subjects. All of the aforementioned blink suppression tasks were completed outside of the scanner, but the same subjects also completed one blink suppression task repeated twice inside of an MRI scanner as well, rating their discomfort only at the end of each completed scan block consisting of 5 suppression trials. We expected that continuous rating during the fMRI trials would change the nature of the task and that this change would be reflected in the BOLD activity. Rather, we will combine the timing of the recorded blinks inside the scanner with the pathophysiological model created for each subject during the suppression tasks outside of the scanner. The degree to which each voxel’s time–signal curve matches the predicted time–discomfort curve, convolved with an individually estimated, model-free 16-s hemodynamic response period, will be computed using a general linear model to create a statistical image of urge-related voxels. The timing of individual tics observed on the video recordings can be entered separately to remove tic-related signal (and analyzed separately for signal related to tics and failure of tic inhibition). Many candidate regions have already been identified by previous studies (*e.g.*, the inferior frontal gyrus, (Yaniv & Lavidor, 2017)), however, we will investigate regions including the supplementary motor area (SMA), anterior cingulate cortex and the caudate. The SMA is a prime region of interest because it is consistently associated with tic generation and control (Ganos, Roessner, & Münchau, 2013) and direct electrical stimulation of the SMA caused the anticipation of movement or the ‘urge’ to execute a movement, without the motor activity itself (Fried et al., 1991). Similarly, fMRI studies by (Wang et al., 2011) have shown that the anterior cingulate cortex and caudate are more active during effective tic suppression. We expect these findings will correlate across subjects with blink suppression success, and possibly with tic suppression success in subjects with TS. This model may allow for a more individualized approach and reflect a more accurate neuronal signal with less noise than would the previous one-size-fits-all models.

## Acknowledgements

The authors thank Dr. Aleksandra Królak of the Department of Medical Electronics, Institute of Electronics, Lodz University of Technology, Łódź, Poland, for kindly sharing her C++ code for eyeblink detection from a live video feed (Królak & Strumiłło, 2012).

## Appendix

## A. Initial data visualization and model construction

In order to investigate the relationship between discomfort intensity and blink occurrence uninterrupted by a blink, we used Python and Matplotlib to plot the discomfort of each subject prior to the first blink in each trial (Hunter, 2007). There was substantial variance within and across subjects, but the central tendency was an increasing curve that appeared to flatten out over time. We initially used a logarithmic functional form, *y*(*t*) = *a*log(*t* + *c*) + *b*, which reasonably described the averaged data from either TS or control subjects. However, the log function initially gave unreliable estimates for the model parameters, so we eventually abandoned it for a delayed square root function,

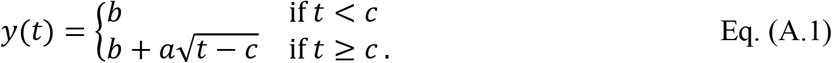

**Figure A.3.**
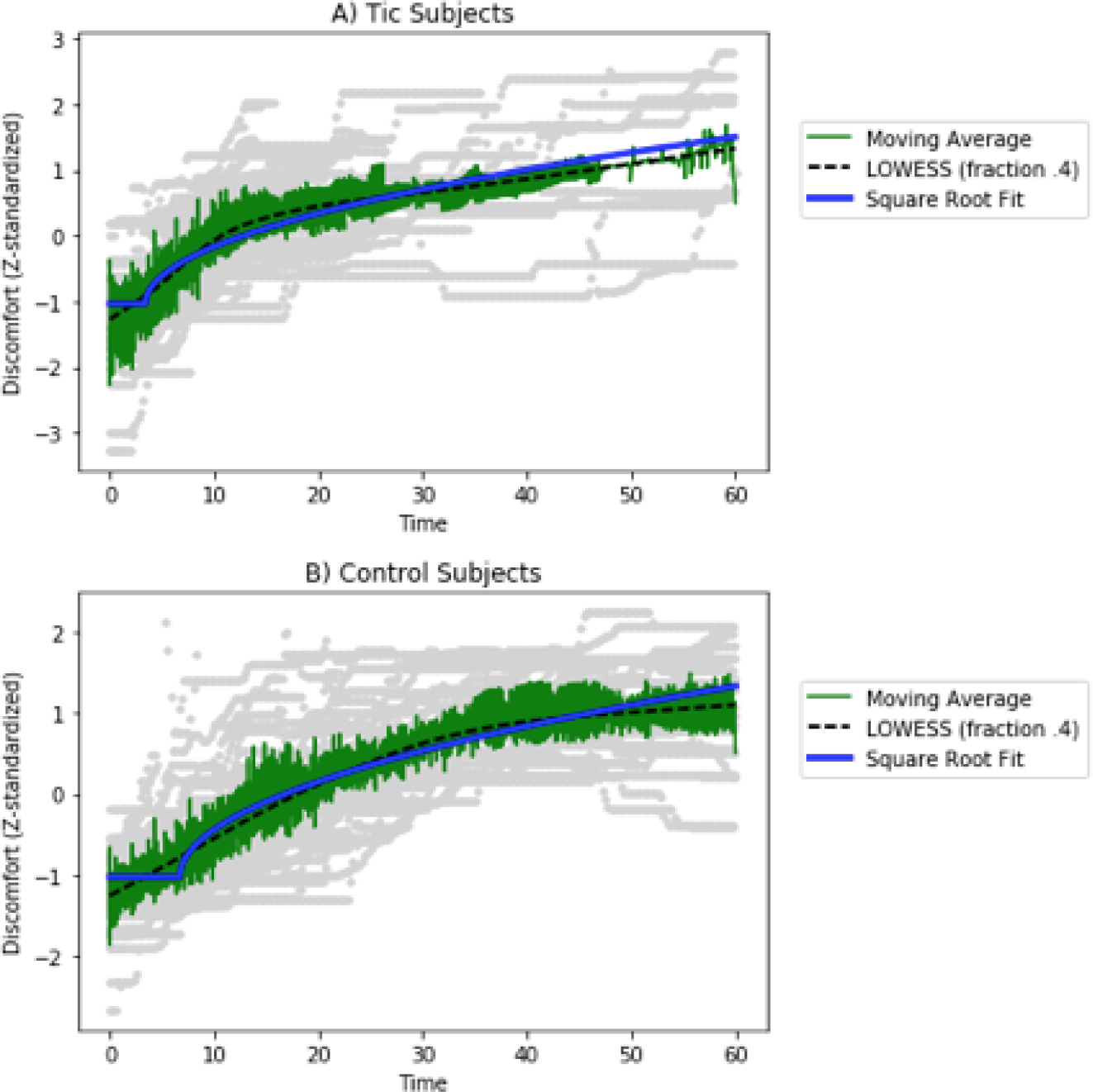
Discomfort during successful blink suppression. Discomfort is plotted across all blink suppression trials, beginning at the start of each “don’t blink” trial and extending to the first blink. The functional form 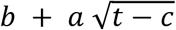 was fit to the data and compared to a LOWESS (locally weighted scatterplot smoothing) fit to the data. Individual subject data is plotted in gray, while the mean is shown in green and LOWESS in black. (A) Data from tic subjects; (B) data from control subjects.

We then wanted to examine the effect of eye closure on discomfort. To do so we plotted the discomfort in between blinks (more than 2 seconds apart) from Study 2. We examined whether the discomfort would dip after a blink and how long it would take to return back to the predicted discomfort. The difference in reported discomfort and expected discomfort (based on the square root fit to the data before the first blink and the observed discomfort when the blink started) was then plotted using the equation

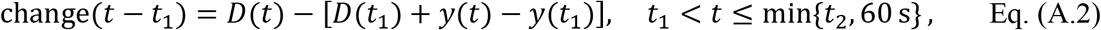

where *D*(*t*) represents the subject’s discomfort ratings at time *t*, *y*(*t*) is the delayed square root function (Eq. A.1) fit to the subject’s data prior to the first blink, *t*_1_ indicates the time point at which the first blink occurred, and *t*_2_ the time at which the second blink occurred. The observed relief in discomfort showed substantial variation, but a tendency to a small, transient dip in discomfort after eye closure, as hypothesized, which we modeled as a parabola passing through (0,0) with vertex at (*h*, *k*), *h* > 0, *k* < 0.

Together, these data and assumptions led to the following model for discomfort during the “don’t-blink” blocks. For a given time point *t*_*i*_ in the interval [0,60s],

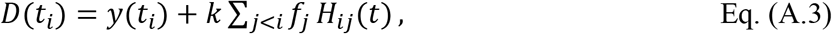

where *y*(*t*) is from Eq. 1, *f*_*j*_ is the fraction of each quarter-second time bin *j* during which the eyes were actually closed, *kH*_*ij*_(*t*) represents the sum of the expected drop in discomfort from each recent 0.25s eye closure,

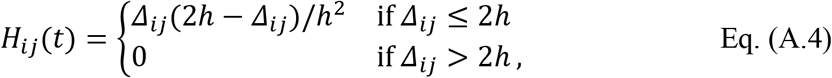

and *Δ*_*ij*_ = *t*_*i*_ − *t*_*j*_. Most of the *f*_*j*_ terms are zero, simplifying the calculation somewhat.

We modeled discomfort during the 30-second “OK to blink” block that immediately followed the 60 seconds of blink suppression based on the mean subject data, which suggested approximately exponential decay, *dD*/*dt* = −*r*(*D* − *b*), with discomfort decay rate constant *r*. We added a factor to account for the presumed modification by eye closure in the preceding time frame, resulting in this difference equation for *t* in the interval (60 s, 90 s):

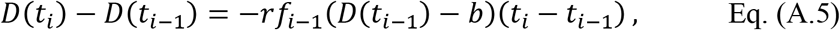

where *f*_*i*−1_ is the fraction of time the subject’s eyes were closed in the previous interval.

Together, Equations A.3 and A.5 form the model we fit to the data.

